# Possible transmission flow of SARS-CoV-2 based on ACE2 features

**DOI:** 10.1101/2020.10.08.332452

**Authors:** Sk. Sarif Hassan, Shinjini Ghosh, Diksha Attrish, Pabitra Pal Choudhury, Vladimir N. Uversky, Bruce D. Uhal, Kenneth Lundstrom, Nima Rezaei, Alaa A. A. Aljabali, Murat Seyran, Damiano Pizzol, Parise Adadi, Antonio Soares, Tarek Mohamed Abd El-Aziz, Ramesh Kandimalla, Murtaza Tambuwala, Gajendra Kumar Azad, Samendra P. Sherchan, Wagner Baetas-da-Cruz, Kazuo Takayama, Ángel Serrano-Aroca, Gaurav Chauhan, Giorgio Palu, Adam M. Brufsky

**Author notes:** Corresponding author *Email addresses* (Sk. Sarif Hassan), (Vladimir N. Uversky), (Kenneth Lundstrom). (Shinjini Ghosh), (Diksha Attrish), (Pabitra Pal Choudhury), (Bruce D. Uhal), (Nima Rezaei), (Alaa A. A. Aljabali), (Murat Seyran), (Damiano Pizzol), (Parise Adadi), (Antonio Soares), (Tarek Mohamed Abd El-Aziz), (Ramesh Kandimalla), (Murtaza Tambuwala), (Gajendra Kumar Azad), (Samendra P. Sherchan), (Wagner Baetas-da-Cruz), (Kazuo Takayama), (Ángel Serrano-Aroca), (Gaurav Chauhan), (Giorgio Palu), (Adam M. Brufsky).

## Abstract

Angiotensin-converting enzyme 2 (ACE2) is the cellular receptor for the Severe Acute Respiratory Syndrome Coronavirus 2 (SARS-CoV-2) that is engendering the severe coronavirus disease 2019 (COVID-19) pandemic. The spike (S) protein receptor-binding domain (RBD) of SARS-CoV-2 binds to the three sub-domains viz. amino acids (aa) 22-42, aa 79-84, and aa 330-393 of ACE2 on human cells to initiate entry. It was reported earlier that the receptor utilization capacity of ACE2 proteins from different species, such as cats, chimpanzees, dogs, and cattle, are different. A comprehensive analysis of ACE2 receptors of nineteen species was carried out in this study, and the findings propose a possible SARS-CoV-2 transmission flow across these nineteen species.

## 1. Introduction

We have been acquainted with the term beta-coronavirus for about two decades when we first encountered the Severe Acute Respiratory Syndrome Coronavirus (SARS-CoV) outbreak that emerged in 2002, infecting about 8000 people with a 10% mortality rate [1]. It was followed by the emergence of Middle East Respiratory Syndrome Coronavirus (MERS-CoV) in 2012 with 2300 cases and mortality rate of 35% [2]. The third outbreak, caused by SARS-CoV-2, was first reported in December 2019 in China, Wuhan province, which rapidly took the form of a pandemic [3, 4]. To date, this new human coronavirus has affected 36 million people worldwide and is held accountable for over one million deaths [5]. SARS-CoV-2 is an enveloped single-stranded plus sense RNA virus whose genome is about 30kb in length, and which encodes for 16 non-structural proteins, four structural and six accessory proteins [6]. The four major structural proteins which play a vital role in viral pathogenesis are Spike protein (S), Nucleocapsid protein (N), Membrane protein (M), and Envelope protein (E) [7, 8]. SARS-CoV-2 infection is mainly characterized by pneumonia [9]; however, multi-organ failure involving myocardial infarction, hepatic, and renal damage is also reported in patients infected with this virus [10]. SARS-CoV-2 binds to the Angiotensin-converting enzyme 2 (ACE2) receptor on the host cell surface via its S protein [11, 12]. ACE2 plays an essential role in viral attachment and entry [13]. The study of the interaction of ACE2 and S protein is of utmost importance [14, 15].

The S1 subunit of the S protein has two domains, the C-terminal and the N-terminal domains, which fold independently, and either of the domains can act as Receptor Binding Domain (RBD) for the interaction and binding to the ACE2 receptor widely expressed on the surface of many cell types of the host [16, 17]. The human ACE2 protein is 805 amino acid long, containing two functional domains: the extracellular N-terminal claw-like peptidase M2 domain and the C-terminal transmembrane collectrin domain with a cytosolic tail [18]. The RBD of the S protein binds to three different regions of ACE2, which are located at amino acids (aa) 24-42, 79-84, and 330-393 positions present in the claw-like peptidase domain of ACE2 [13]. ACE2 modulates angiotensin activities, which promote aldosterone release and increase blood pressure and inflammation, thus causing damage to blood vessel linings and various types of tissue injury [19]. ACE2 converts Angiotensin II to other molecules and reduces this effect [20]. However, when SARS-CoV-2 binds to ACE2, the function of ACE2 is inhibited and in turn leads to endocytosis of the virus particle into the host cell [21].

Zoonotic transmission of this virus from bat to human and random mutations acquired by SARS-CoV-2 during human to human transmission has also empowered this virus with the ability to undergo interspecies transmission and recently many cases have been reported stating that different species can be infected by this virus [22].

In this study, we aim to determine the susceptibility of other species, whether they bear the capability of being a possible host of SARS-CoV-2. We chose nineteen different species and analyzed the ACE2 protein sequence in relation to the human ACE2 sequence and determined the degree of variability by which the sequences differed from each other. We performed a comprehensive bioinformatics analysis in addition to the phylogenetic analysis based on full-length sequence homology, polarity along with individual domain sequence homology and secondary structure prediction of these protein sequences. These finding could be emerged to six distinct clusters of nineteen species based on the collective analysis and thereby provided a prediction of the interspecies SARS-CoV-2 transmission.

## 2. Data and Methods

### 2.1. Data acquisition and findings

The ACE2 protein receptor sequences from nineteen species *Homo sapiens* (Human), *Capra hircus* (Domestic goat), *Pan troglodytes* (Chimpanzee), *Equus caballus* (Horse), *Salmo salar* (Atlantic salmon), *Mesocricetus auratus* (Golden hamster), *Rhinolophus ferrumequinum* (Greater horseshoe bat), *Pteropus alecto* (Black flying fox), *Mustela putorius furo* (Domestic ferret), *Danio rerio* (Zebrafish), *Manis javanica* (Sunda pangolin), *Sus scrofa* (Domestic pig), *Macaca mulatta* (Rhesus macaque), *Bos taurus* (Aurochs), *Pelodiscus sinensis* (Chinese soft-shelled turtle), *Pteropus vampyrus* (Large flying fox), *Rattus norvegicus* (Brown rat), *Felis catus* (Domestic cat), and *Gallus gallus* (Red jungle fowl) were derived from the NCBI database [23]. Nineteen species and their respective ACE2 protein accession IDs with length are presented in Table-1.

**Table 1:** Nineteen species and their associated ACE2 sequences

| Name | Species | ACE2 Accession ID | Length |
| --- | --- | --- | --- |
| S1 | *Homo sapiens* | NP_001358344.1 | 805 |
| S2 | *Pan troglodytes* | PNI38577.1 | 805 |
| S3 | *Macaca mulatta* | XP_028697658.1 | 805 |
| S4 | *Equus caballus* | XP_001490241.1 | 805 |
| S5 | *Felis catus* | NP_001034545.1 | 805 |
| S6 | *Mesocricetus auratus* | XP_005074266.1 | 805 |
| S7 | *Manis javanica* | XP_017505752.1 | 805 |
| S8 | *Pteropus alecto* | XP_006911709.1 | 805 |
| S9 | *Capra hircus* | AHI85757.1 | 804 |
| S10 | *Sus scrofa* | NP_001116542.1 | 805 |
| S11 | *Mustela putorius furo* | XP_004758943.1 | 805 |
| S12 | *Bos taurus* | NP_001019673.2 | 804 |
| S13 | *Rattus norvegicus* | NP_001012006.1 | 805 |
| S14 | *Pteropus vampyrus* | XP_011361275.1 | 804 |
| S15 | *Rhinolophus ferrumequinum* | XP_032963186.1 | 805 |
| S16 | *Gallus gallus* | XP_416822.2 | 808 |
| S17 | *Pelodiscus sinensis* | XP_006122891.1 | 808 |
| S18 | *Danio rerio* | XP_005169417.1 | 807 |
| S19 | *Salmo salar* | XP_014062928.1 | 695 |

The nearest neighborhood phylogeny of the nineteen species derived from the NCBI public server based on ACE2 protein sequence similarity is shown in Fig.1 (A) [24].

**Figure 1:**
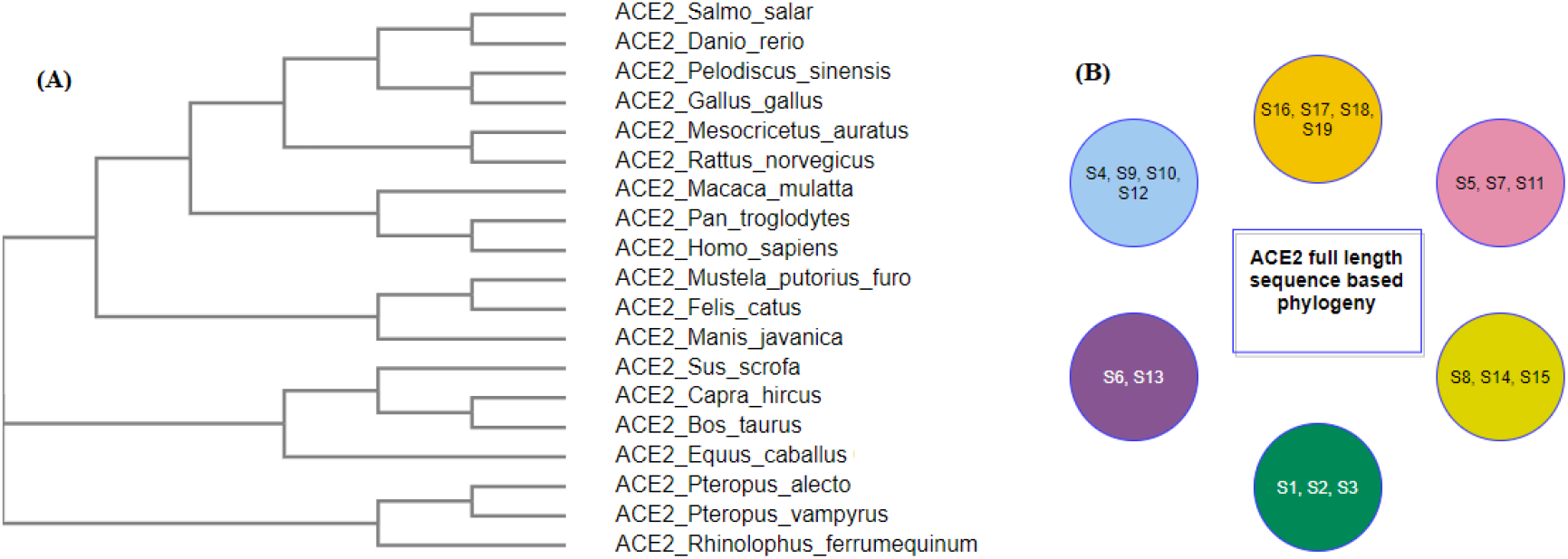
ACE2 full-length sequence-based phylogeny among nineteen species (A) and its derived clusters (B)

ACE2 sequence similarity among the species derives six clusters as shown in (Fig.1 (B)). The contact residues of the receptor-binding domain (RBD) of the spike protein (YP_009724390.1) of SARS-CoV-2 with the homo sapiens ACE2 interface are presented in Table-2 [13].

**Table 2:** Contact residues of RBD spike protein of SARS-CoV-2 and Homo sapiens ACE2

| SARS-CoV-2 RBD | ACE2 (Homo sapiens) | SARS-CoV-2 RBD | ACE2 (Homo sapiens) |
| --- | --- | --- | --- |
| K417 | Q24 | Q493 | Q42 |
| G446 | T27 | G496 | L79 |
| Y449 | F28 | Q498 | M82 |
| Y453 | D30 | T500 | Y83 |
| L455 | K31 | N501 | N330 |
| F456 | H34 | G502 | K353 |
| A475 | E35 | Y505 | G354 |
| F486 | E37 |  | D355 |
| N487 | D38 |  | R357 |
| Y489 | Y41 |  | R393 |

The three designated domains, D1 (aa 24-42), D2 (aa 79-84) and D3(aa 330-393) respectively contain the residues which bind to the RBD of the S protein.

### 2.2. Methods

#### Examining amino acid substitutions

For human ACE2 receptor, substitutions were examined for all species and only those substitutions are accounted for, which occurred in the binding residues in the mentioned three domains D1, D2 and D3 [13]. Based character of the substitutions which interfered with the binding residues of the ACE2 across various species, two types were defined: substitutions affected transmission (M1), and substitutions did not affect transmission (M2).

Multiple sequence alignments and associated phylogenetic trees were developed using the NCBI web-suite across all the individual binding domains D1, D2, and D3 of ACE2 of the nineteen species [25, 26].

#### K-means clustering

Algorithmic clustering technique derives homogeneous subclasses within the data such that data points in each cluster are as similar as possible according to a widely used distance measure viz. Euclidean distance. One of the most commonly used simple clustering techniques is the *K-means clustering* [27, 28]. The algorithm is described below in brief:

##### Algorithm

K-means algorithm is an iterative algorithm that tries to form equivalence classes from the feature vectors into K (pre-defined) clusters where each data point belongs to only one cluster [27].

- Assign the number of desired clusters (*K*) (in the present study, *K* = 6).
- Find centroids by first shuffling the dataset and then randomly selecting *K* data points for the centroids without replacement.
- Keep iterating until there is no change to the centroids.
- Find the sum of the squared distance between data points and all centroids.
- Assign each data point to the closest cluster (centroid).
- Compute the centroids for the clusters by taking the average of the all data points that belong to each cluster.

In this present study, nineteen species were clustered using *Matlab* by inputting the distance matrix derived from the feature vectors associated with the three domains of ACE2 across all species.

#### Secondary structure predictions

The secondary structure of full-length ACE2 sequence of all species were predicted using the web-server CFSSP (Chou & Fasman Secondary Structure Prediction Server) [29]. This server predicts secondary structure regions from the protein sequence such as alpha-helix, beta-sheet, and turns from the amino acid sequence [29]. On obtaining the full-length ACE2 secondary structures, individual domains D1, D2, and D3 were cropped for each species.

#### Bioinformatics features

Several bioinformatics features viz. Shannon entropy, instability index, aliphatic index, charged residues, half-life, melting temperature, N-terminal of the sequence, molecular weight, extinction coefficient, net charge at pH7 and isoelectric point of D1, D2, and D3 domains of ACE2 for all nineteen species were determined using the web-servers *Pfeature and ProtParam* [30, 31].

##### Shannon entropy

It measures the amount of complexity in a primary sequence of ACE2. It was determined using the web-server *Pfeature* by the formula

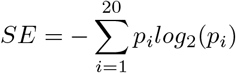

where *p_i_* denotes the frequency probability of a given amino acid in the sequence [30].

*Instability index* is determined using the web-server *ProtParam* and it estimates the stability of a protein in a test tube. A protein whose instability index is smaller than 40 is predicted as stable. A value above 40 predicts that the protein may be unstable [30].

*Aliphatic index* of a protein is defined as the relative volume gathered by aliphatic side chains (alanine, valine, isoleucine, and leucine). It may be regarded as a positive factor for increasing the thermostability of globular proteins, such as ACE2 [30].

##### N-terminal

It was reported that the N-terminal of a protein is responsible for its function. For each domain sequence, N-terminal residue was determined using the *Pfeature* [30].

##### In vivo half-life

The half-life predicts the time it takes for half of the protein amount to degrade after its synthesis in the cell. The N-end rule originated from the observations that the identity of the N-terminal residue of a protein plays an essential role in determining its stability in vivo [32].

##### Extinction coefficients

The extinction coefficient measures how much light a protein absorbs at a particular wavelength. It is useful to estimate this coefficient when a protein is getting purified [32].

#### Polarity sequence

Every amino acid in the domains D1, D2 and D3 of ACE2 were recognized as polar (P) and non-polar (Q) and thus every D1, D2 and D3 for all the species turned out to be binary sequences with two symbols P and Q. Then homology of these sequences for each domain was made and consequently, phylogenetic relationship was drawn.

## 3. Results

Based on amino acid homology, secondary structures, bioinformatics and polarity of the three domains D1, D2 and D3 of ACE2, all nineteen species were clustered. Finally a cumulative set of nineteen species clusters was built, among which the SARS-CoV-2 transmission may occur.

### 3.1. Phylogeny and clustering based on ACE2 domain-based homology

First, we examined all the substitutions with similar properties and similar side chain binding atoms, signifying that the substitutions would not impede the SARS-CoV-2 transmission. Note that, all the mutations are considered concerning the human ACE2 domains D1, D2 and D3 (Fig.2).

**Figure 2:**
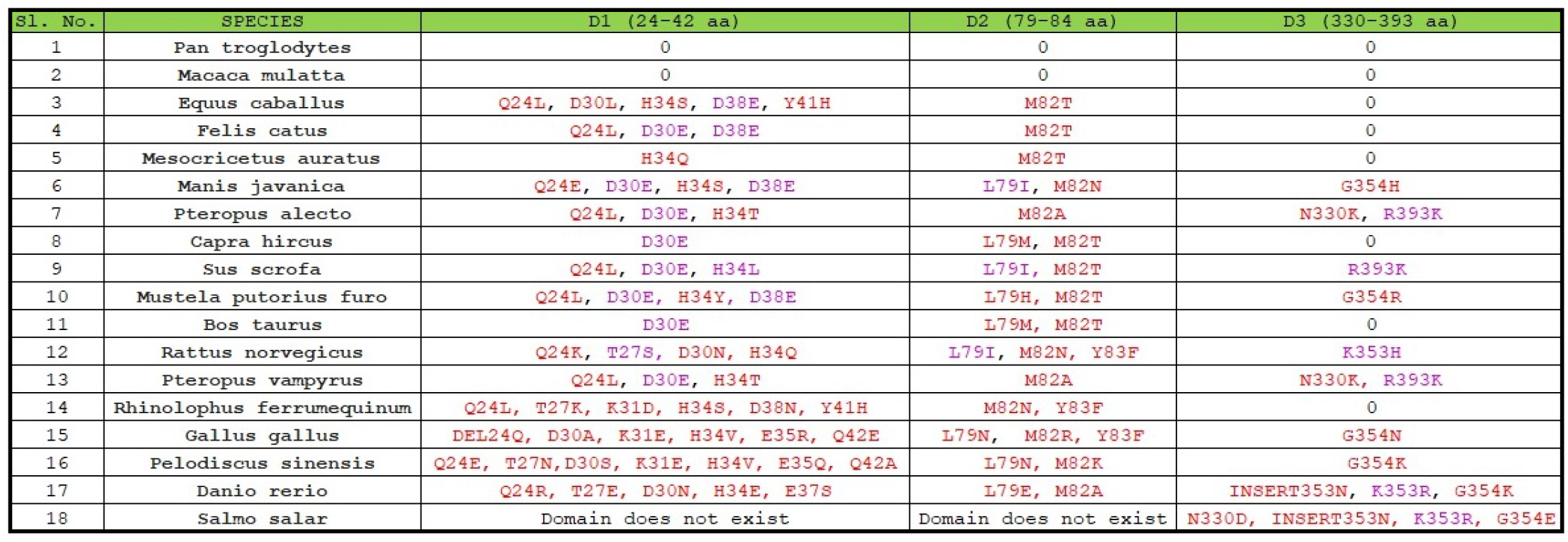
Substitutions in D1, D2 and D3 domains of ACE2 across eighteen species

### In D1 domain

out of eighteen species, eight species were found to possess a substitution at position 30 where D (Aspartate) was substituted by E (Glutamate) and four species were found to carry the D38E substitution. It was reported that in the aspartate side chain, the oxygen atom was involved in ionic-ionic interaction and the side-chain oxygen atom was also present in glutamate, so this substitution may not affect the protein-protein interaction properties [33, 34]. In the T27S substitution, Threonine and Serine both possess OH that participates in binding and in the H34L substitution, both Histidine and Leucine use the NH group for interaction with another amino acid. Consequently, if we consider only the critical perspective for these substitutions, we can conclude that these changes would not impede the binding between the S and ACE2 protein.

### In D2 domain

L79I bears importance across eighteen species since both these amino acids (Leucine and isoleucine) share similar chemical properties. So, if we analyze the changes in amino acid residues based on their chemical properties, which is the main contributing factor for protein-protein interaction, we can conclude that it will not significantly affect the binding between ACE-2 and RBD of the S protein.

### In D3 domain

out of eleven substitutions, three substitutions (R393K, K353H, and K353R) were observed of the similar type with similar side chain interacting atoms and therefore changes at these positions would not affect the interaction of ACE2 with that of the S protein.

Secondly, across all nineteen species, homology were derived based on amino acid sequences, and consequently, associated phylogenetic trees were drawn (Fig.3).

**Figure 3:**
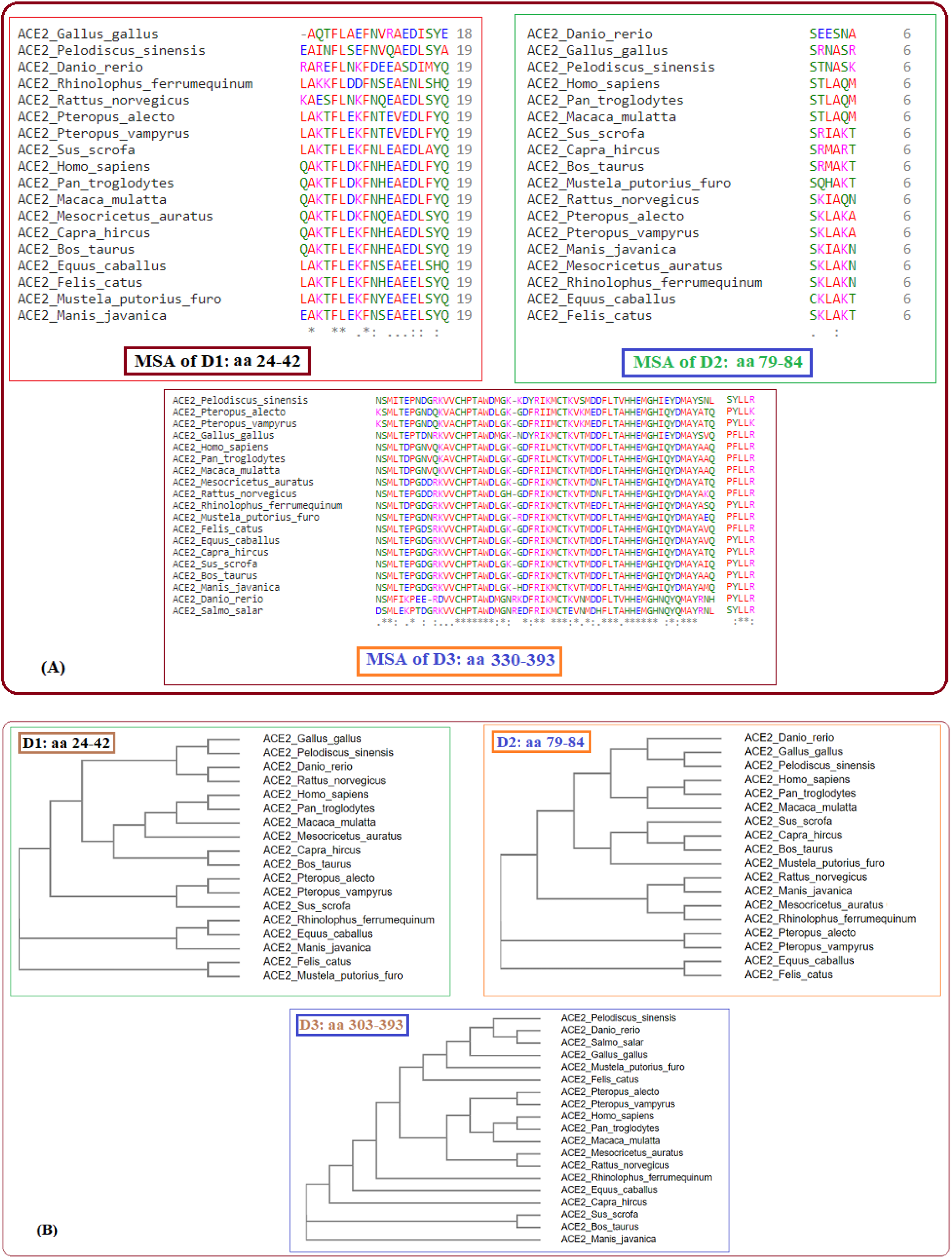
Multiple sequence alignments of D1, D2 and D3 domains of ACE2 of nineteen species (A) and respective phylogenies (B).

Six clusters of the nineteen species were formed using the K-means clustering technique based on sequence homology of the three domains (Fig 4). The clusters of species {*S*1, *S*2, *S*3} and {*S*6, *S*13} stayed together for the ACE2 full-length sequence homology and the combination of three domain-based sequence similarity. The species S16, S17, and S18 also followed the same as observed.

**Figure 4:**
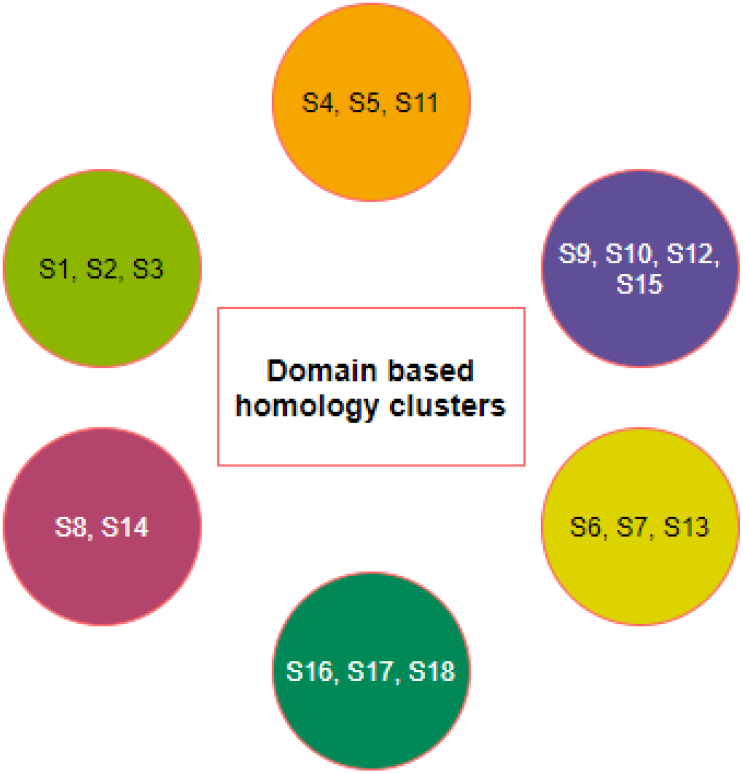
Clusters of species based on domain-based sequence homology

Further, it was observed that sequence homology of the D1, D2, and D3 domains clustered the species S15 into the cluster where S9, S10 and S12 belong, although S15 was similar to the ACE2 sequence of S8 and S9. Although the species S4 was very much similar to S9, S10, and S12 for full-length ACE2 homology, it clubbed with S5 and S11 concerning the three domain-based sequence spatial organizations. Also S7 was found to be staying in the proximity of S6 and S13 although S7 was very much similar to S5 and S11 based on ACE2 homology.

### 3.2. Clustering based on secondary structures

For each domain of ACE2 of nineteen species, the secondary structure was predicted (Fig.5). For each domain species are grouped into several subgroups.

**Figure 5:**
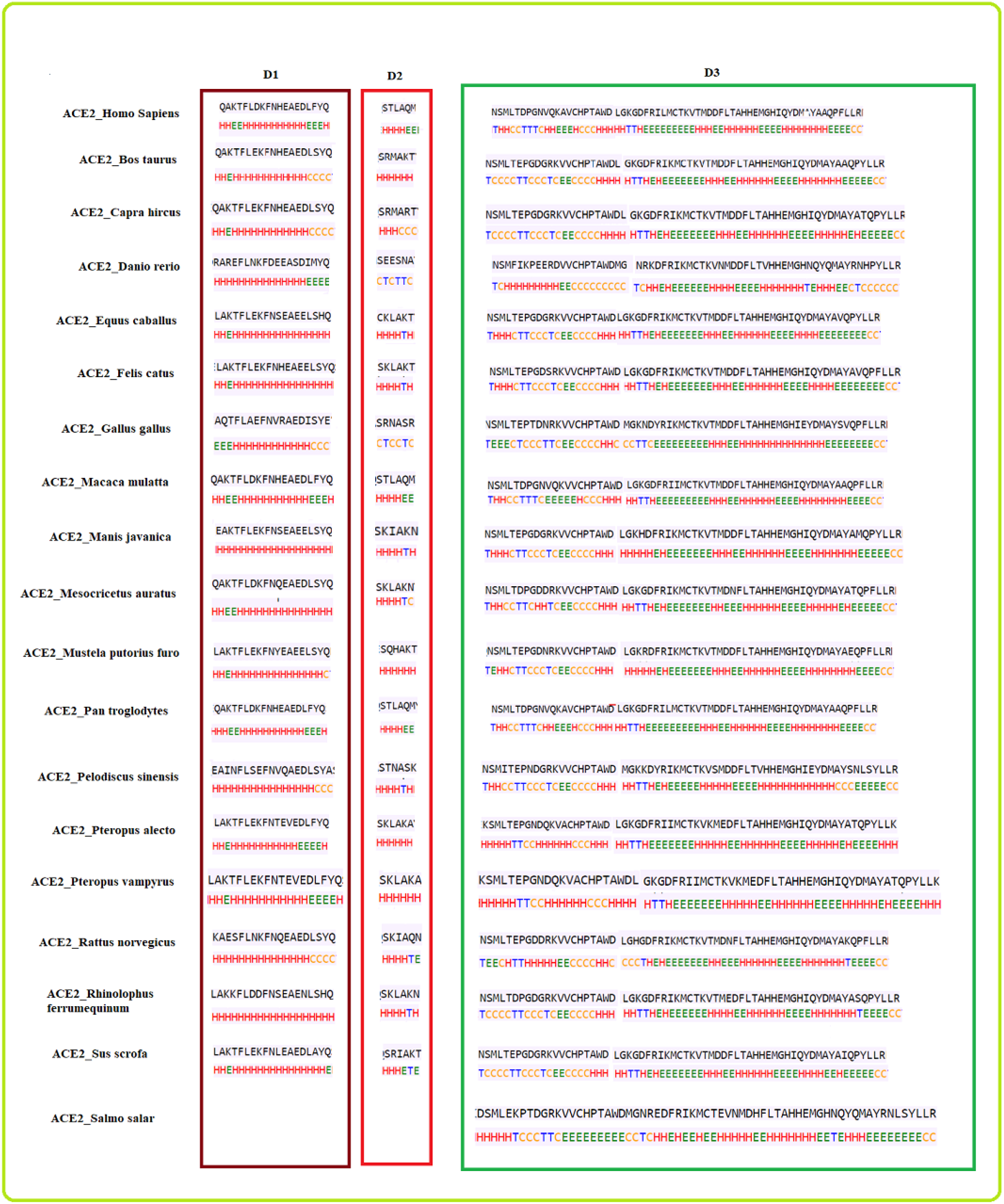
Predicted secondary structures of D1, D2, and D3 domains for all the species

#### With respect to the D1 domain

- *Bos taurus* and *Capra hircus*
- *Equus caballus* and *Felis catus*
- *Mustela putorius furo* has a structure closer to *Equus caballus* and *Felis catus* as it has only one difference of a coil present at position 42. Also, *Mesocricetus auratus* has a secondary structure similar to the above two except an extended helix at position four. Similarly, *Sus scrofa* which has an extended sheet instead of a helix at position 42. So, *Mustela putorius furo, Mesocricetus auratus*, and *Sus scrofa* can be put in the same cluster as *Equus caballus* and *Felis catus*.
- *Homo sapiens, Macaca mulatta* and *Pan troglodytes*
- *Manis javanica* and *Rhinolophus ferrumequinum*
- *Pteropus alecto* and *Pteropus Vampyrus*
- *Rattus norvegicus* and *Pelodiscus sinensis* have similar structures differing by the presence of an extra coil at position 39 for *Rattus norvegicus*.
- *Gallus gallus* and *Danio rerio* has a unique secondary structure in comparison to the others.

These individual six clusters show six different secondary structures in D1 shared by sixteen species, which shows high similarities in their secondary structures, while the rest two have a unique secondary structure for D1 domain. Thus, these six clusters have similar secondary structures indicating that the species in the six clusters are closely related.

#### With respect to the D2 domain

- *Homo sapiens, Macaca mulatta* and *Pan troglodytes*
- *Bos taurus, Mustela putorius furo, Pteropus alecto* and *Pteropus vampyrus*
- *Equus caballus, Felis catus, Manis javanica, Pelodiscus sinensis* and *Rhinolophus ferrumequinum*
- *Danio rerio* and *Gallus gallus*

Similarly, for D2 domain, we found four clusters with the same secondary structure, indicating that they are closely related.

#### With respect to the D3 domain

- *Homo sapiens* and *Pan troglodytes*
- *Bos Taurus, Rhinolophus ferrumequinum, Sus scrofa* and *Capra hircus*
- *Equus caballus* and *Felis catus*
- *Pteropus alecto* and *Pteropus vampyrus*

Again for D3 domain, four different clusters were bearing similar secondary structures therefore these species are also closely related.

Based on the similarity among the three domains, all nineteen species were clustered (Fig.6).

**Figure 6:**
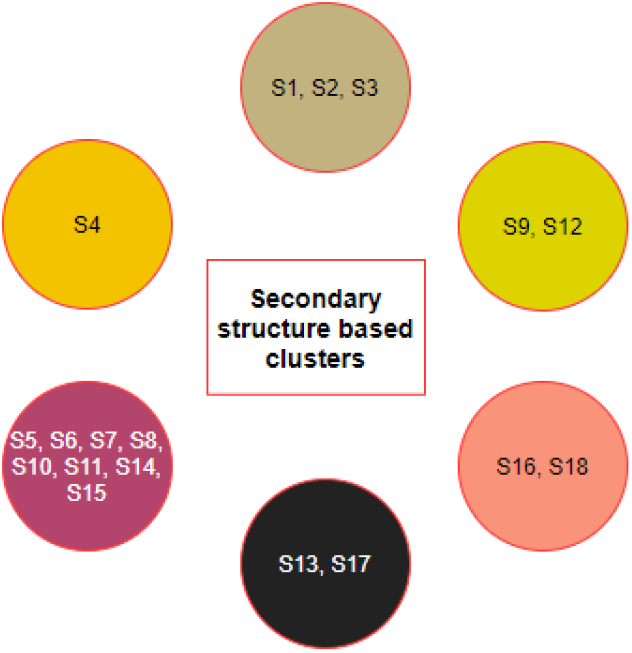
Clusters of species based on the secondary structures of the D1, D2 and D3 domains

From the clusters (Fig.6) based on the secondary structure of the three domains of ACE2, it was observed that the species S4 was clustered uniquely, though S4 is clustered with S9 and S19 based on ACE2 full-length sequence homology. Furthermore, S6 and S13 were found to be similar based on ACE2 homology, but they got clustered into two different clusters when the secondary structure of three domains was concerned. In contrast, the group of species {*S*1,*S*2,*S*3}, {*S*9,*S*12}, {*S*16, *S*18}, and {*S*5, *S*7*S*11} remained in the same clusters concerning to ACE2 homology as well as individual secondary structures of the domains.

### 3.3. Clustering based on bioinformatics

Twelve bioinformatics features viz. Shannon entropy, Instability Index, aliphatic index, charged residues, half-life, melting temperature, N-terminal of the sequence, molecular weight, extinction coefficient, net charge at pH7, and isoelectric point of the D1, D2, and D3 domains of ACE2 for all nineteen species were determined (Fig.7).

**Figure 7:**
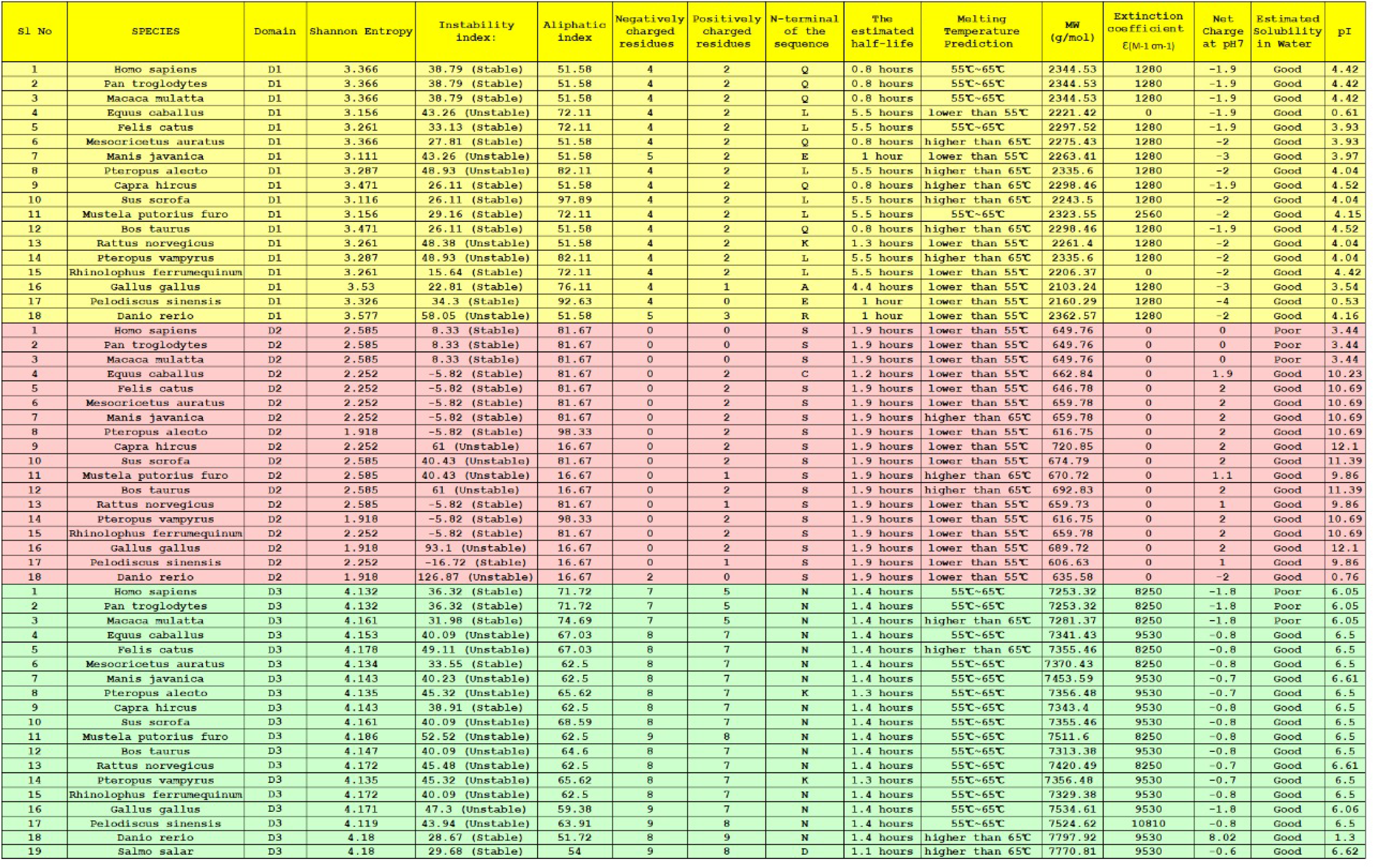
Bioinformatics of the D1, D2, and D3 domains of ACE2 from nineteen species

For each species, a twelve-dimensional feature vector was found (Fig.7). For each domain D1, D2, and D3 domain, distance matrix was determined using the Euclidean distance

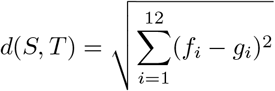

. Note that here *f_i_* and *g_i_* denote the *i*th feature for the species *S* and *T* respectively. This distance matrices with heatmap representation for all three domains are presented in Figs. 8, 9 & 10. In addition, by inputting the distance matrix, using the K-means clustering technique, several clusters of species were formed for each domain (Figs. 8, 9 and 10).

**Figure 8:**
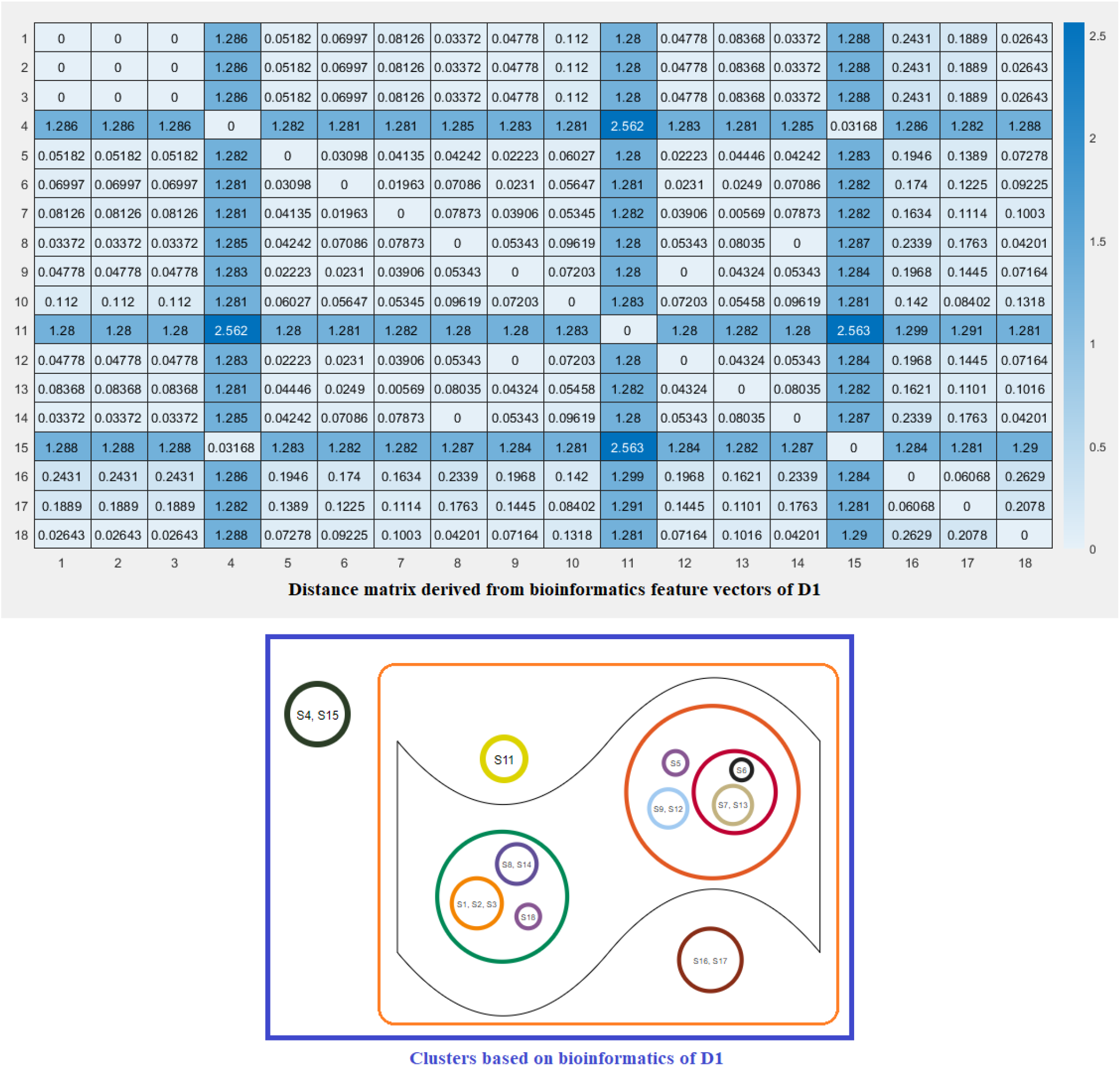
Distance matrix based on the bioinformatics feature vectors of D1 of ACE2 across eighteen species and associated clusters

**Figure 9:**
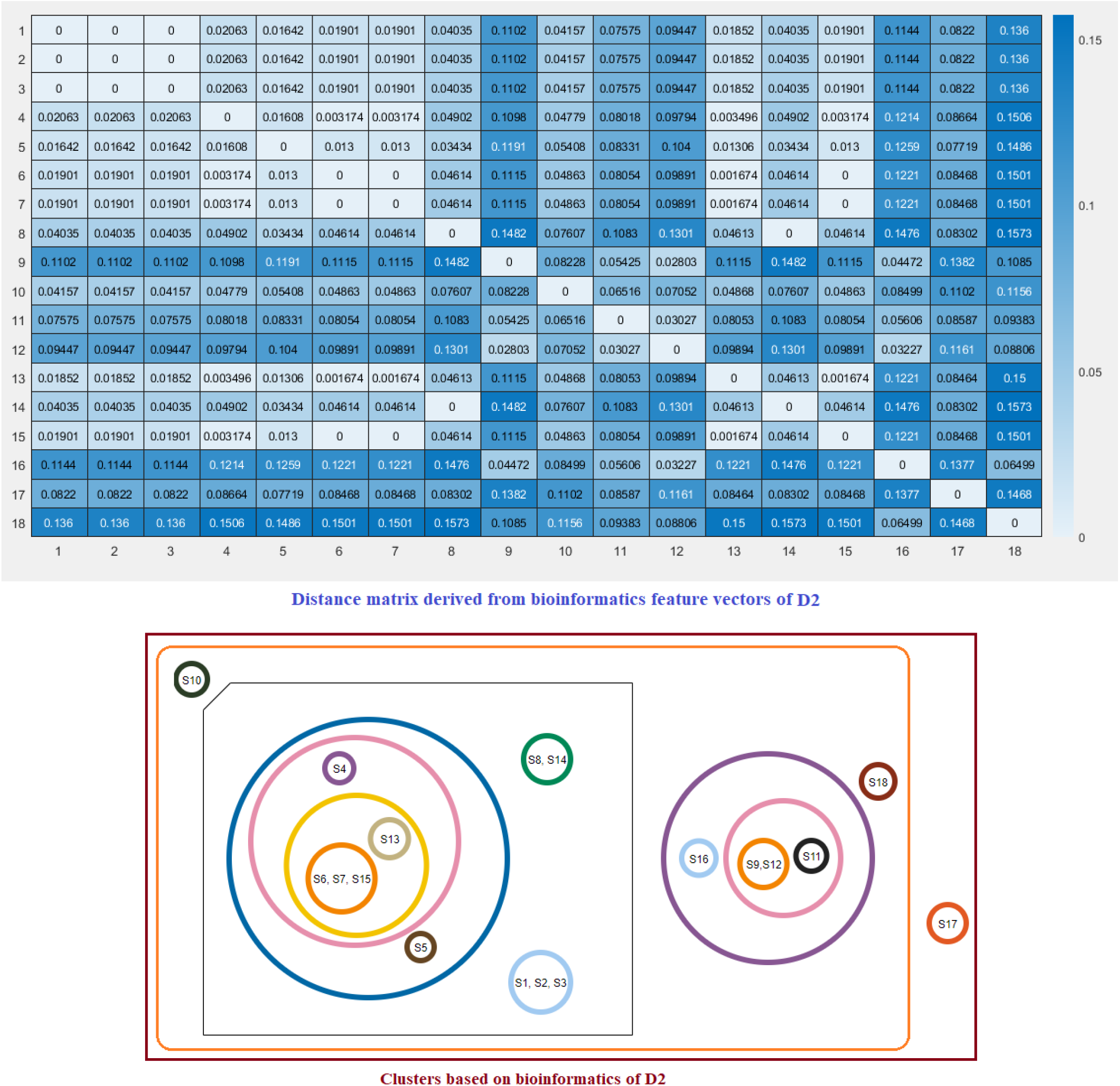
Distance matrix based on the bioinformatics feature vectors of D2 of ACE2 across eighteen species and associated clusters

**Figure 10:**
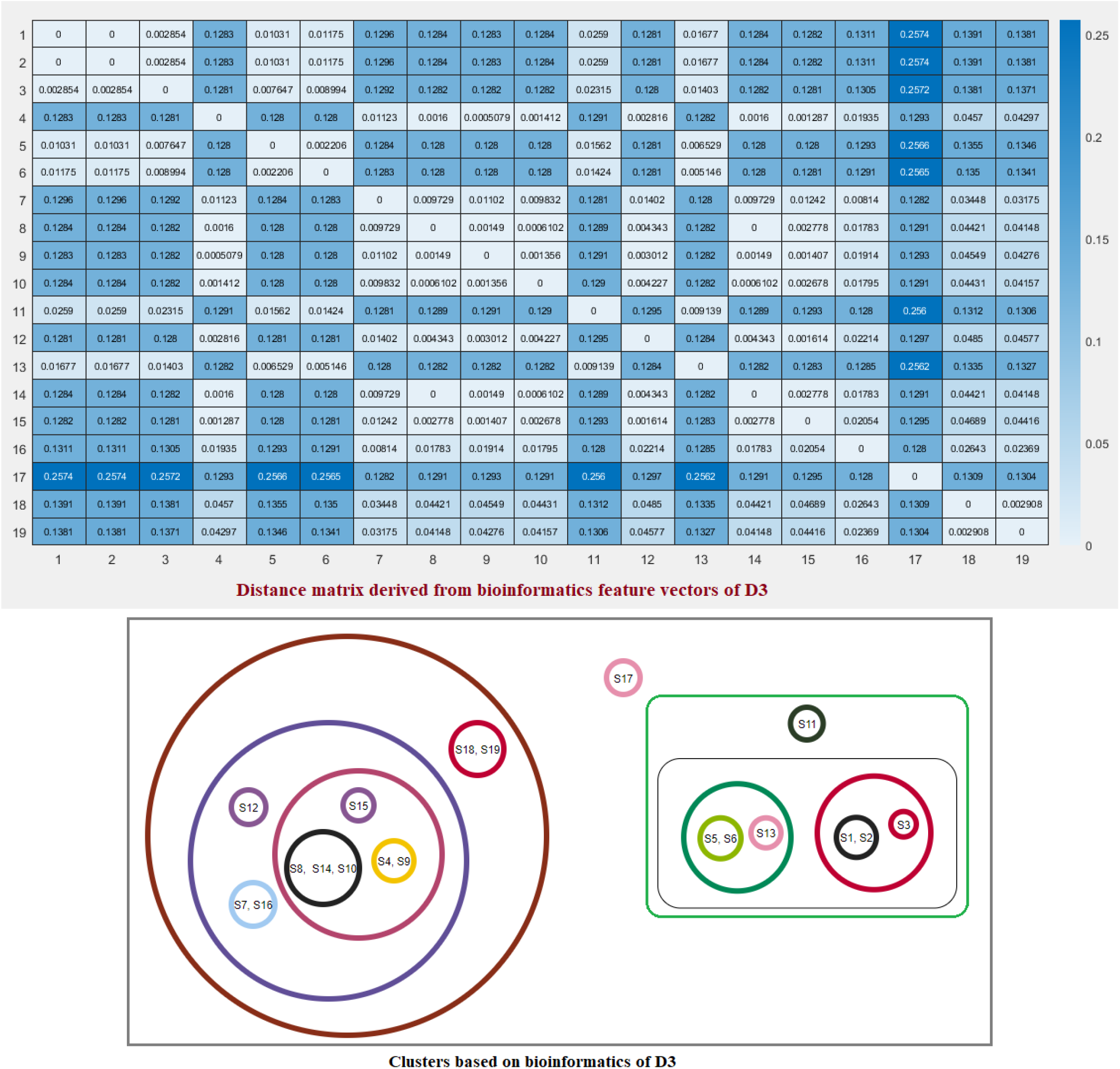
Distance matrix based on the bioinformatics feature vectors of D3 of ACE2 across nineteen species and associated clusters

A final set of six clusters was formed using the K-means clustering method to have all three domains for the nineteen different species (Fig. 11). Although the species S7 was clustered with the species S5 and S11 as per full-length ACE2 sequence homology, S7 formed a unique singleton cluster when the bioinformatics features were taken into consideration. Similarly, the species S16 formed a singleton cluster though it was clustered with S17, S18, and S19 as per amino acid homology of ACE2. The sequence homology of ACE2 made the four species S16, S17, S18, and S19 into a single cluster but bioinformatics features placed the species S18 in a cluster where the other three species S1, S2, and S3 belonged. Based on bioinformatics features, S4 clustered together with S15 though the ACE2 receptor of S4 was sequentially similar to ACE2 of S9, S10, and S12.

**Figure 11:**
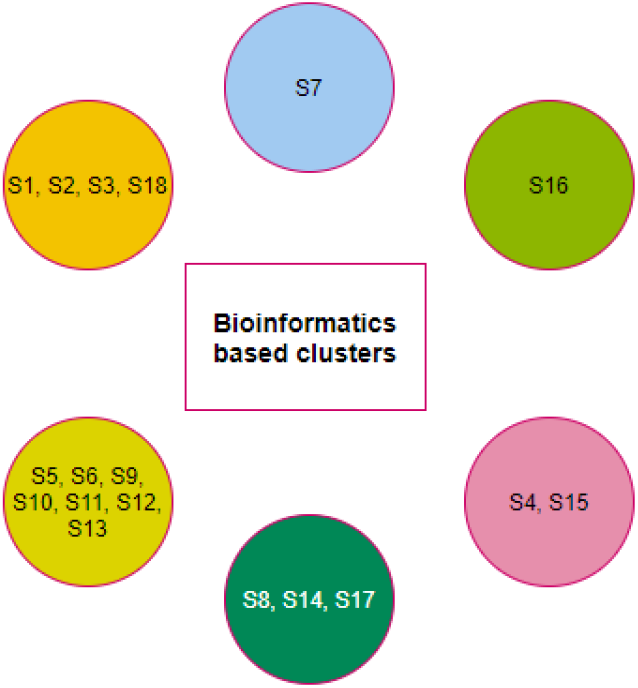
Clusters of species based on the bioinformatics of the D1, D2, and D3 domains

The clusters {*S*1, *S*2, *S*3}, {*S*6, *S*13}, and {*S*9, *S*10, *S*12} were unaltered with respect to the full length ACE2 homology and bioinformatics features.

### 3.4. Phylogeny and clustering based on polarity

In the D1 domain, it was observed that across eighteen species, polarity of thirteen amino acids among nineteen (24-42 aa) amino acids were found to be conserved. Based on the polarity and non-polarity nature of the amino acids, the species were arranged in a phylogenetic tree (Fig. 12).

**Figure 12:**
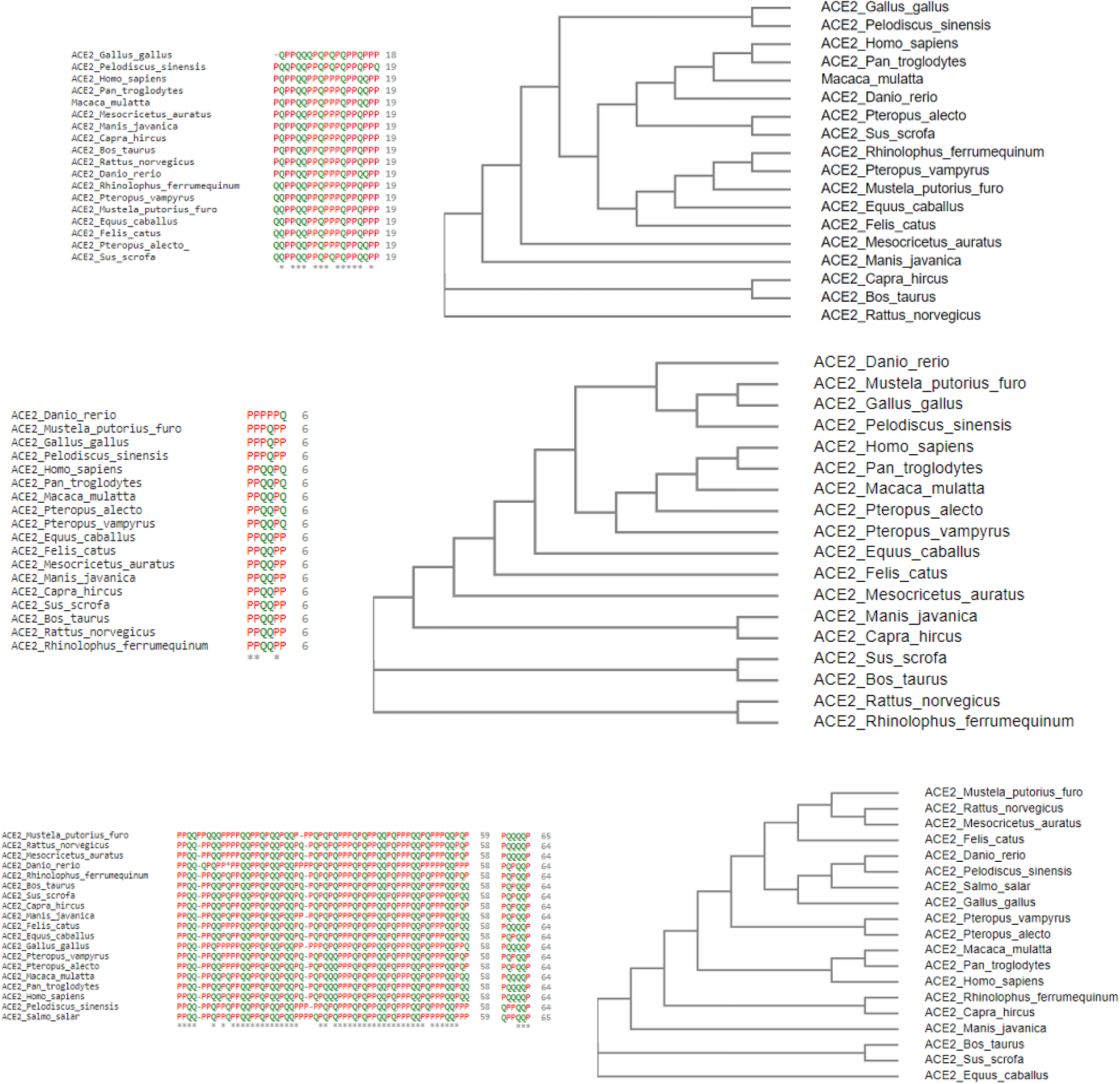
Polarity sequence of the D1, D2 and D3 domains across all species alignment and associated phylogenetic relationships

It was found that *Homo sapiens, Pan troglodyte, Macaca mulatta* and *Danio rerio* were closer according to this analysis. *Pteropus alecto, Pteropus vampyrus* and *Sus scrofa* were occurred in parallel along with the above three and formed a different clade indicating the closeness based on polarity. Again, the case for *Gallus gallus, Rhinolophus ferrumequinum, Mustela putorius furo, Equus caballus* and *Felis catus* is similar. Two separate groups, *Mesocricetus auratus, Manis javanica* and *Capra hircus, Bos taurus*, were similarly placed nearby, indicating that the polarity of amino acids of the proteins for these species was similar. *Pelodiscus sinensis* and *Rattus norvegicus* occurred separately and were not grouped with any other species but bears similarity with both the groups containing species *Mesocricetus auratus, Manis javanica and Pteropus alecto, Pteropus vampyrus, Sus scrofa*.

In the D2 domain, out of six amino acid long sequences, the polarity of three amino acids was conserved across 18 species, and among them, one amino acid was a binding residue. *Homo sapiens, Pan troglodytes, Macaca mulatta, Pteropus vampyrus* and *Pteropus alecto* were grouped together since the overall polarity of their amino acid chain was found to be similar and simultaneously, *Danio rerio, Mustela putorius furo, Gallus gallus* and *Pelodiscus sinensis* were placed together. Also, three groups comprising *Manis javanica, Capra hircus* & *Bos taurus, Sus scrofa* & *Rattus norvegicus, Rhinolophus ferrumequinum* respectively were placed in close proximity based on their polarity and non-polarity of the amino acids in the protein sequence. However, *Equus caballus, Felis cattus*, and *Mesocricetus auratus* were placed separately since they did not show much resemblance based on polarity.

In the D3 domain of ACE2 sequences of *Salmo salar* and *Danio rerio* there was an insertion of a polar amino acid in one of the binding residue positions that may affect the binding of ACE2 to that of RBD of SARS-CoV2 negatively. A total of three binding residues were already reported in the D3 domain, of which one of them remained conserved concerning polarity across the nineteen species. *Rattus norvegicus, Mustela putorius furo, Mesocricetus auratus* and *Felis catus* were grouped under a single clade based on the polarity of their protein sequence. Similar was the case for *Danio rerio, Pelodiscus sinensis, Salmo salar*, and *Gallus gallus*. Due to the sequence similarity between *Pteropus vampyrus* and *Pteropus alecto*, their polarity of the protein sequence was also similar and thus grouped. Sequence similarity was also observed for *Homo sapiens, Pan troglodytes* and *Macaca mulatta*, so again these were categorized together. Two groups comprising of *Rhinolophus ferrumequinum*, *Capra hircus* and *Bos taurus, Sus scrofa* respectively, were sorted together indicating their similar nature of polarity and non-polarity of protein sequence. Lastly, *Manis javanica* and *Equus caballus* were placed separately signifying that the sequence of both the species were quite distinct.

The individual groups of species based on the polarity of individual D1, D2 and D3 domains have emerged into six disjoint clusters (Fig. 13).

**Figure 13:**
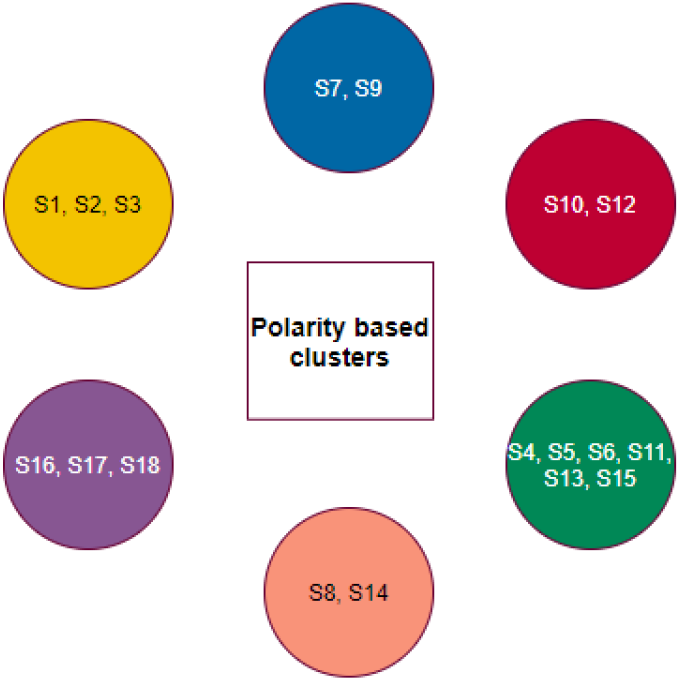
Clusters based on groups of species based on domain-wise polarity

Here the clusters {*S*1, *S*2, *S*3}, {*S*16, *S*17, *S*18}, {*S*8, *S*14}, {*S*6, *S*13}, and {*S*10, *S*12} remained invariant with regards to the homology of full length ACE2 as well as polarity sequence of the D1, D2 and D3 domains.

### 3.5. Possible clusters of transmission of SARS-CoV-2

Based on all the different clustered formed on the basis of amino acid homology, secondary structures, bioinformatics, and polarity of the D1, D2, and D3 domains of ACE2, final clusters of all nineteen species were devised using the K-means clustering method Fig. 14.

**Figure 14:**
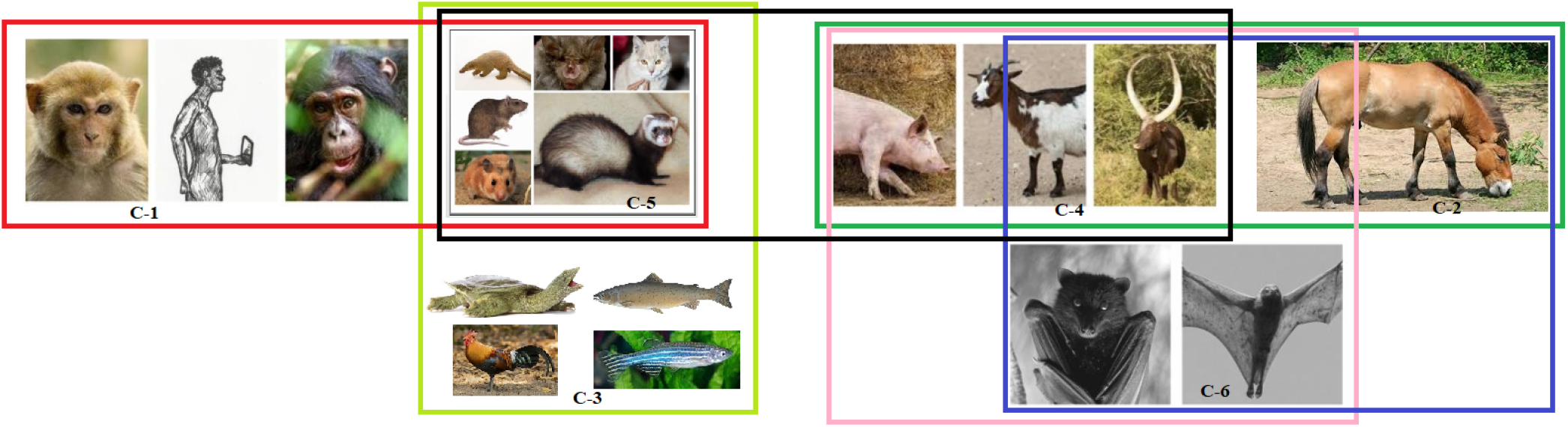
Schematic representation of a possible set of clusters of transmission of SARS-CoV-2

In Fig. 14, it was found that the cluster-1 (C-1) comprising of *Homo sapiens, Pan troglodyte*, and *Macaca mulatta* were close to cluster-5 (C-5) comprising of *Felis catus* (Cat), *Mesocricetus auratus* (Golden Hamster), *Manis javanica* (Sunda pangolin), *Mustela putorius furo* (Ferret), *Rattus norvegicus* (Rat), and *Rhinolphus ferrumequinum* (Greater horseshoe bat). This C-5 is also close to cluster-3 (C-3) [*Gallus gallus* (red jungle fowl), *Pelodiscus sinensis* (Chinese shell turtle), *Danio rerio* (zebrafish) and *Salmo salar*], and cluster-4 (C-4) [*Capra hircus* (Goat), *Bos taurus* (Cattle) and *Sus scrofa* (pig)]. C-4 also showed resemblance with cluster-2 (C-2) [*Pteropus alecto*, and *Pteropus vampyrus*], and cluster-6 (C-6) that comprised of *Equus caballus* (horse) only. Although, both C-2 and C-6 were also close to each other.

Furthermore, a pooled analyses based on the two type of substitutions (one is affecting SARS-CoV-2 transmission (M1), and the other one is SARS-CoV-2 non-affecting transmission(M2)) for all the six final clusters are presented in Table-3.

**Table 3:**
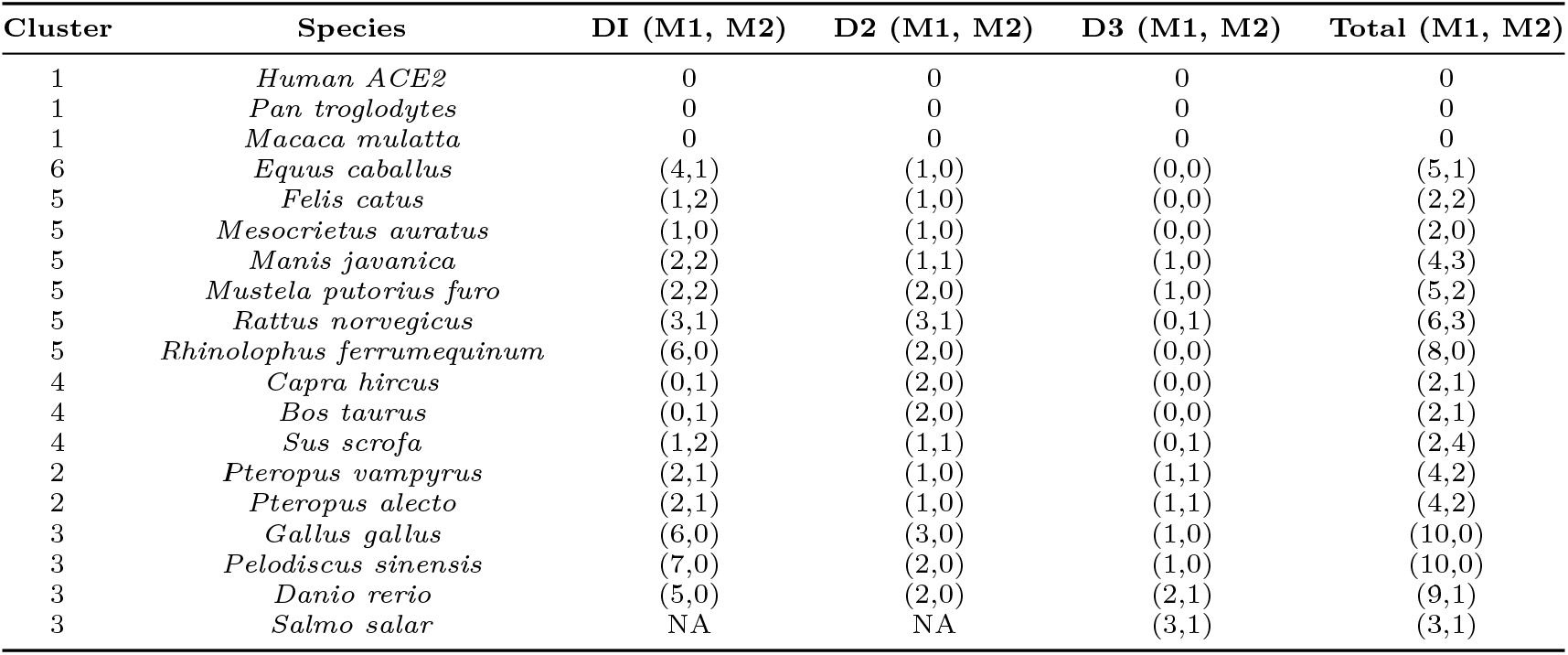
M1 and M2 substitutions across nineteen ACE2 receptors

| Cluster | Species | D1 (M1, M2) | D2 (M1, M2) | D3 (M1, M2) | Total (M1, M2) |
| --- | --- | --- | --- | --- | --- |
| 1 | *Human ACE2* | 0 | 0 | 0 | 0 |
| 1 | *Pan troglodytes* | 0 | 0 | 0 | 0 |
| 1 | *Macaca mulatta* | 0 | 0 | 0 | 0 |
| 6 | *Equus caballus* | (4,1) | (1,0) | (0,0) | (5,1) |
| 5 | *Felis catus* | (1,2) | (1,0) | (0,0) | (2,2) |
| 5 | *Mesocrietus auratus* | (1,0) | (1,0) | (0,0) | (2,0) |
| 5 | *Manis javanica* | (2,2) | (1,1) | (1,0) | (4,3) |
| 5 | *Mustela putorius furo* | (2,2) | (2,0) | (1,0) | (5,2) |
| 5 | *Rattus norvegicus* | (3,1) | (3,1) | (0,1) | (6,3) |
| 5 | *Rhinolophus ferrumequinum* | (6,0) | (2,0) | (0,0) | (8,0) |
| 4 | *Capra hircus* | (0,1) | (2,0) | (0,0) | (2,1) |
| 4 | *Bos taurus* | (0,1) | (2,0) | (0,0) | (2,1) |
| 4 | *Sus scrofa* | (1,2) | (1,1) | (0,1) | (2,4) |
| 2 | *Pteropus vampyrus* | (2,1) | (1,0) | (1,1) | (4,2) |
| 2 | *Pteropus alecto* | (2,1) | (1,0) | (1,1) | (4,2) |
| 3 | *Gallus gallus* | (6,0) | (3,0) | (1,0) | (10,0) |
| 3 | *Pelodiscus sinensis* | (7,0) | (2,0) | (1,0) | (10,0) |
| 3 | *Danio rerio* | (5,0) | (2,0) | (2,1) | (9,1) |
| 3 | *Salmo salar* | NA | NA | (3,1) | (3,1) |

Based on the Table-3 information regarding the number of M1 and M2 substitutions, the intra-species transmission of SARS-CoV-2 were enlightened as follow:

- C-1: None of the species bear any mutation in the binding residues and are conserved, so viral transmission is immaculate.
- C-2: This cluster has an equal number of transmission affecting and transmission non-affecting types of substitutions. Therefore, both have an equal probability of getting infected from each other.
- C-3: Here again, *Gallus gallus, Pelodiscus sinensis*, and *Danio rerio* have a similar ratio of S1 to S2, signifying possible flow of viral transmission within these three species. However, *Salmo salar* is unique and distant, and therefore, the probability of viral transmission is unlikely.
- C-4: The species in this cluster have a similar number of transmission-affecting and transmission non-affecting type of substitutions which shows that the flow of viral transmission would be continuous among these three species.
- C-5: Transmission between *Felis catus* and *Mesocrietus auratus* is highly likely which is the same for *Manis javanica, Mustela putorius furo*, and *Rattus norvegicus* as indicated by their similar number of substitutions. Therefore, the inter-transmission between these species is highly plausible. While *Rhinolophus ferrumequinum* has a relatively high value of transmission affecting substitutions from all of the above, its susceptibility to getting infected from others species is uncertain.
- C-6: A total of five transmission affecting substitutions in the three domains for human were observed.

## 4. Discussion

In this study, we amassed the ACE2 protein sequences of nineteen species to investigate the possible transmission of SARS-CoV-2 among these species in relation to human ACE2 protein. Multiple sequence alignments of these ACE2 receptors enabled us to estimate the similarity concerning amino acids and from that, we observed that *Salmo salar* (Salmon fish) was quite distant. It also gave us the idea that some of the amino acid substitutions in the binding residues occurring across the species with respect to human ACE2 were resulted in amino acids having similar binding properties, indicating that their interactions with RBD of the S protein will be similar to that of humans, thus making transmission across these species feasible. It was observed that *Homo sapiens* and *Pan troglodytes* (Chimpanzee) have complete sequence similarity. In contrast, *Macaca mulatta* (Rhesus macaque) shared a high percentage of sequence identity except for two amino acid positions. However, no substitutions were observed in the binding of amino acid residues, making the viral transmission across these species highly likely. Again, *Pteropus vampyrus* (Large flying fox) and *Pteropus alecto* (Black flying fox) have precisely the same ACE2 sequence and thus signifying high viral transmission and that both of them have an equal chance of getting infected with each other.

Further analysis led us to present a possible transmission flow among the nineteen species, as illustrated in Fig. 14. The multifaceted examination of the ACE2 protein indicated that interspecies SARS-CoV-2 transmission is quite possible and we have tried to provide a better insight into it by predicting the possible transmission among species within the same cluster and between clusters too. However, further in-depth analysis is necessary in the future for the identification of new hosts of SARS-CoV-2 as well as for determination of possible ways to prevent inter-species transmission.

## Author Contributions

SSH conceived the problem. SSH, DA, SG carried out the work. SSH analyzed the results and wrote the primary draft of the article. All authors reviewed, edited, and approved the final manuscript.

## References

[1] W. H. Organization, Severe acute respiratory syndrome (sars), Weekly Epidemiological Record 78 (13) (2003) 89–89.

[2] W. H. Organization, Outbreak news: severe acute respiratory syndrome (sars), Weekly Epidemiological Record 78 (12) (2003) 81–83.

[3] J.-j. Zhang, X. Dong, Y.-y. Cao, Y.-d. Yuan, Y.-b. Yang, Y.-q. Yan, C. A. Akdis, Y.-d. Gao, Clinical characteristics of 140 patients infected with sars-cov-2 in wuhan, china, Allergy (2020).

[4] Y. Liu, Z. Ning, Y. Chen, M. Guo, Y. Liu, N. K. Gali, L. Sun, Y. Duan, J. Cai, D. Westerdahl, et al., Aerodynamic analysis of sars-cov-2 in two wuhan hospitals, Nature 582 (7813) (2020) 557–560.

[5] W. H. Organization, et al., Coronavirus disease 2019 (covid-19): situation report, 82 (2020).

[6] Y.-Z. Zhang, E. C. Holmes, A genomic perspective on the origin and emergence of sars-cov-2, Cell (2020).

[7] M. Zheng, L. Song, Novel antibody epitopes dominate the antigenicity of spike glycoprotein in sars-cov-2 compared to sars-cov, Cellular & molecular immunology 17 (5) (2020) 536–538.

[8] R. Minakshi, A. T. Jan, S. Rahman, J. Kim, A testimony of the surgent sars-cov-2 in the immunological panorama of the human host, Frontiers in Cellular and Infection Microbiology 10 (2020) 539.

[9] X. Yang, Y. Yu, J. Xu, H. Shu, H. Liu, Y. Wu, L. Zhang, Z. Yu, M. Fang, T. Yu, et al., Clinical course and outcomes of critically ill patients with sars-cov-2 pneumonia in wuhan, china: a single-centered, retrospective, observational study, The Lancet Respiratory Medicine (2020).

[10] S. Zaim, J. H. Chong, V. Sankaranarayanan, A. Harky, Covid-19 and multi-organ response, Current Problems in Cardiology (2020) 100618.

[11] F. Saponaro, G. Rutigliano, S. Sestito, L. Bandini, B. Storti, R. Bizzarri, R. Zucchi, Ace2 in the era of sars-cov-2: controversies and novel perspectives (2020).

[12] S. Li, J. Han, A. Zhang, Y. Han, M. Chen, Z. Liu, M. Shao, W. Cao, Exploring the demographics and clinical characteristics related to the expression of angiotensin-converting enzyme 2, a receptor of sars-cov-2, Frontiers in Medicine 7 (2020) 530.

[13] J. Lan, J. Ge, J. Yu, S. Shan, H. Zhou, S. Fan, Q. Zhang, X. Shi, Q. Wang, L. Zhang, et al., Structure of the sars-cov-2 spike receptor-binding domain bound to the ace2 receptor, Nature 581 (7807) (2020) 215–220.

[14] M. Hussain, N. Jabeen, F. Raza, S. Shabbir, A. A. Baig, A. Amanullah, B. Aziz, Structural variations in human ace2 may influence its binding with sars-cov-2 spike protein, Journal of medical virology (2020).

[15] P. McMillan, B. D. Uhal, Covid-19–a theory of autoimmunity to ace-2, MOJ immunology 7 (1) (2020) 17.

[16] L. Samavati, B. D. Uhal, Ace2, much more than just a receptor for sars-cov-2, Frontiers in Cellular and Infection Microbiology 10 (2020) 317.

[17] G. K. Veeramachaneni, V. Thunuguntla, J. Bobbillapati, J. S. Bondili, Structural and simulation analysis of hotspot residues interactions of sars-cov 2 with human ace2 receptor, Journal of Biomolecular Structure and Dynamics (2020) 1–11.

[18] W. Li, C. Zhang, J. Sui, J. H. Kuhn, M. J. Moore, S. Luo, S.-K. Wong, I.-C. Huang, K. Xu, N. Vasilieva, et al., Receptor and viral determinants of sars-coronavirus adaptation to human ace2, The EMBO journal 24 (8) (2005) 1634–1643.

[19] M. Gheblawi, K. Wang, A. Viveiros, Q. Nguyen, J.-C. Zhong, A. J. Turner, M. K. Raizada, M. B. Grant, G. Y. Oudit, Angiotensin-converting enzyme 2: Sars-cov-2 receptor and regulator of the renin-angiotensin system: celebrating the 20th anniversary of the discovery of ace2, Circulation research 126 (10) (2020) 1456–1474.

[20] C. Tikellis, M. Thomas, Angiotensin-converting enzyme 2 (ace2) is a key modulator of the renin angiotensin system in health and disease, International journal of peptides 2012 (2012).

[21] K. Gorshkov, K. Susumu, J. Chen, M. Xu, M. Pradhan, W. Zhu, X. Hu, J. C. Breger, M. Wolak, E. Oh, Quantum dot-conjugated sars-cov-2 spike pseudo-virions enable tracking of angiotensin converting enzyme 2 binding and endocytosis, ACS nano (2020).

[22] V. N. Uversky, F. Elrashdy, A. Aljadawi, E. M. Redwan, Household pets and sars-cov2 transmissibility in the light of the ace2 intrinsic disorder status, Journal of Biomolecular Structure and Dynamics (2020) 1–4.

[23] K. D. Pruitt, T. Tatusova, D. R. Maglott, Ncbi reference sequence (refseq): a curated non-redundant sequence database of genomes, transcripts and proteins, Nucleic acids research 33 (suppl_1) (2005) D501–D504.

[24] S. Federhen, The ncbi taxonomy database, Nucleic acids research 40 (D1) (2012) D136–D143.

[25] M. Johnson, I. Zaretskaya, Y. Raytselis, Y. Merezhuk, S. McGinnis, T. L. Madden, Ncbi blast: a better web interface, Nucleic acids research 36 (suppl_2) (2008) W5–W9.

[26] J. P. Jenuth, The ncbi, in: Bioinformatics Methods and Protocols, Springer, 2000, pp. 301–312.

[27] A. Likas, N. Vlassis, J. J. Verbeek, The global k-means clustering algorithm, Pattern recognition 36 (2) (2003) 451–461.

[28] Y. Lu, S. Lu, F. Fotouhi, Y. Deng, S. J. Brown, Fgka: A fast genetic k-means clustering algorithm, in: Proceedings of the 2004 ACM symposium on Applied computing, 2004, pp. 622–623.

[29] T. A. Kumar, Cfssp: Chou and fasman secondary structure prediction server, Wide Spectrum 1 (9) (2013) 15–19.

[30] A. Pande, S. Patiyal, A. Lathwal, C. Arora, D. Kaur, A. Dhall, G. Mishra, H. Kaur, N. Sharma, S. Jain, et al., Computing wide range of protein/peptide features from their sequence and structure, bioRxiv (2019) 599126.

[31] V. K. Garg, H. Avashthi, A. Tiwari, P. A. Jain, P. W. Ramkete, A. M. Kayastha, V. K. Singh, Mfppi–multi fasta protparam interface, Bioinformation 12 (2) (2016) 74.

[32] C. N. Pace, F. Vajdos, L. Fee, G. Grimsley, T. Gray, How to measure and predict the molar absorption coefficient of a protein, Protein science 4 (11) (1995) 2411–2423.

[33] H.-X. Zhou, X. Pang, Electrostatic interactions in protein structure, folding, binding, and condensation, Chemical reviews 118 (4) (2018) 1691–1741.

[34] M. M. Flocco, S. L. Mowbray, Strange bedfellows: interactions between acidic side-chains in proteins, Journal of molecular biology 254 (1) (1995) 96–105.

